# Knock down of Calcineurin-B2, a calcium binding regulatory subunit of Calcineurin, gives rise to hypercontraction myopathy in indirect flight muscles of *Drosophila* through dysregulation of calcium homeostasis

**DOI:** 10.1101/108605

**Authors:** Ruchi Jhonsa, Upendra Nongthomba

## Abstract

Muscle is a calcium responsive tissue and requires calcium for initiation of muscle contraction. Calcium concentration in muscle is tightly regulated by calcium binding proteins. Calcineurin-B2 (canB2), a calcium binding, regulatory subunit of calcineurin, is the isoform maximally expressed in the indirect flight muscles (IFMs) of *Drosophila*. The developmental and functional roles of CanB2 particularly in the maintenance of calcium homeostasis are not understood. In this study, we show that the loss of *canB2* causes hypercontraction of IFMs. Genetic interaction studies with hypercontraction mutants suggest a synergistic interaction between Calcineurin-B2 and structural protein Troponin-T. Similarly, interaction studies with *dSERCA* mutant reveal that Calcineurin-B2 is important for regulating calcium homeostasis in muscles. *In vivo* calcium imaging showed that Calcineurin-B2 deficiency leads to arrhythmicity in the calcium oscillations. We show that Calcineurin-B2 knockdown causes deregulation of calcium homeostasis, which results in unregulated acto-myosin interaction. However, overexpression of Calcineurin-A, which inhibits calcium oscillations, had no effect on myofibrillogenesis suggesting that calcium activation is dispensable for myofibrillar assembly. Our findings contribute to the understanding of muscle physiology in normal as well as pathophysiological conditions.

## Introduction

Cardiac hypertrophy results from a wide variety of stimuli, including hypertension, mechanical overload, ischemia or mutations in genes encoding structural proteins (Marian and Roberts, 1995; Barry et al., 2008; Frey et al., 2004). Despite the diverse range of stimuli that cause cardiac hypertrophy, there is a defined response of cardiac tissue to these signals that includes an increase in protein synthesis followed by increase in cell size, enhanced sarcomeric organization, increased expression of fetal cardiac genes, and immediate-early genes, such as c-*fos* and c-*myc* (reviewed in Chien et al., 1993; Sadoshima and Izumo, 1997). Although the causes of cardiac hypertrophy have been extensively studied, the underlying molecular mechanisms remain poorly understood.

Numerous studies have emphasized the role of deregulated intracellular calcium (Ca^2+^) as a signal for cardiac hypertrophy. Intracellular Ca^2+^ concentration rises in response to muscle stretching or increased loads on working heart preparations (Marban et al., 1987; Bustamante et al., 1991; Hongo et al., 1995), showing the importance of calcium for connecting signals to the physiological response. Several humoral factors, which include angiotensin II (AngII), phenylephrine (PE), and endothelin-1 (ET-1), also increase calcium concentrations in the muscles and are known to induce the hypertrophic response in cardiomyocytes (Sadoshima and Izumo, 1993; Leite et al., 1994; Karliner et al., 1990).

In B and T cells, the calcium-dependent phosphatase, calcineurin has been shown to activate the immune response as a result of an increase in calcium concentration. Calcineurin plays a crucial role in heart morphogenesis and cardiac hypertrophy (de la Pompa et al., 1998, Lim and Molkentin, 2000). Overexpression of an activated form of calcineurin during mouse heart development leads to pronounced cardiac hypertrophy (Molkentin et al., 1998) and inhibition of calcineurin by cyclosporine A and FK-506, can block this hypertrophic response (Sussman et al., 1998). Calcineurin activity is essential for skeletal muscle differentiation and suppression of the activity by cyclosporin A inhibits differentiation *in vitro* (Abbott et al., 1998). Studies have shown that the hypertrophic skeletal muscles have higher expression of calcineurin and it can, in turn, induce the GATA transcription factor, which associates with NFATc1 in the regulation of muscle gene transcription (Musaro et al., 1999; Semsarian et al., 1999). Calcineurin has also been shown to modulate the activity of Mef2 in skeletal muscles and neurons by dephosphorylation of specific serine residues (Wu et al., 2001). In order to identify potential regulators of calcineurin, Sullivan and Rubin (2002) performed a dominant modifier screen by overexpressing an activated form of fly calcineurin in the *Drosophila* eye. The study identified nine complementation groups that either enhanced or suppressed the eye phenotype induced by overexpression of *canA*^*act*^ and identified Egf signaling as one of the pathways regulated by calcineurin (Sullivan and Rubin, 2002). Of the several suppressors identified, *canB2* gene, the regulator of Calcineurin-A phosphatase activity was found to suppress the lethal phenotype induced by Calcineurin-A overexpression. A reduction in the levels of *canB2* led to muscle degeneration via hypercontraction suggesting a role for *canB2* in maintaining the levels of myosin heavy chain protein in *Drosophila* indirect flight muscles (IFMs) (Gajewski et al., 2003, 2006).

Unlike vertebrates, *Drosophila* genome codes for single NFAT transcription factor which lacks calcineurin-binding site. Considering the demonstrated role of calcineurin in several myogenic events in vertebrates and invertebrates, we used the indirect flight muscles (IFMs) of the *Drosophila*, which are physiologically similar to cardiac muscles and structurally similar to skeletal muscles (Peckham et al., 1990; Moore, 2006), to understand how calcineurin may be involved in the pathogenesis of muscle hypertrophy, particularly in the events that are independent of NFAT pathway. Our study shows that the loss of either of the Calcineurin-A isoforms (CanA1 and Pp2B-14D) in the IFMs does not lead to structural or functional impairments, suggesting functional redundancy between these two Calcineurin-A isoforms. In this paper, we show how a reduction in CanB2, the major calcium dependent regulator of CanA in IFMs, leads to hypercontraction myopathy condition by deregulation of acto-myosin interaction and calcium homeostasis.

## Results

### Calcineurin subunits have redundant function in the IFMs

In order to assess the role of Calcineurin A in muscle development and function, all the three isoforms of Calcineurin-A were independently knocked down in IFMs. We did not get sufficient knockdown using *CanA1-14F* RNAi line and hence this line was not used for further experiments. *canA1* and *Pp2B-14D* were knocked down in muscles using *Dmef2*-Gal4 and knockdown was confirmed by semi-quantitative RT-PCR (Fig. 1A,A’). Flies with reduced levels of either *canA1* or *Pp2B-14D* in muscles did not show defects in the flying ability and majority of flies were up-flighted (Fig. 1B,B’). Correlating well with the flight test data, there was no discernible defect in the structure of the IFMs of these flies. Flies with reduced expression of *canA1* or *Pp2B-14D* have normal fascicle organization (Fig. 1D’,D”) and the sarcomere organization of the myofibrils was comparable to the control (Fig. 1D).

**Fig.1.**
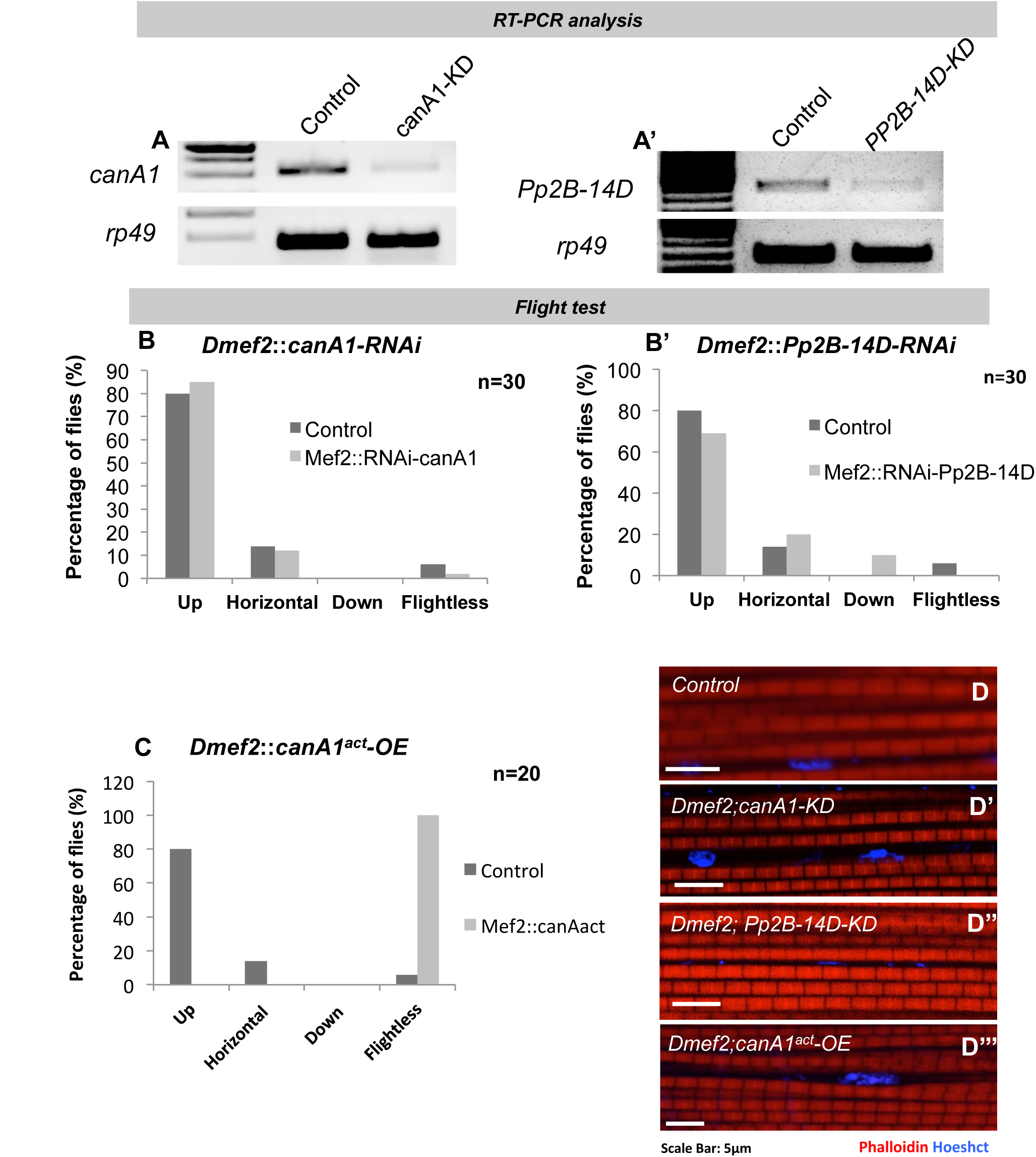
Calcineurin-A isoforms show redundancy in the function. (A) RT-PCR showing *canA1* expression. First lane shows IFM *canA1* expression in wild-type and second lane shows *canA1* expression in *canA1* deficient IFMs (*canA1-RNAi; Dmef2-Gal4*). (A’) RT-PCR showing *Pp2B-14D* expression. First lane shows IFM *Pp2B-14D* expression in wild-type and second lane shows *Pp2B-14D* expression in *Pp2B-14D* deficient IFMs (*Pp2B-14D-RNAi; Dmef2-Gal4;*). (B and B’) Flight test shows that adult *CanA1* and *Pp2B-14D* deficient flies are largely flighted (n=30). (C) Flight test shows that all the flies overexpressing *canA1* are flightless (n=30). (D,D’,D”,D”’) (D) Wild type adult, (D’) *canA1* deficient flies (*canA1*-RNAi; *Dmef2*-*Gal4*), (D’’) *Pp2B-14D* deficient flies (*Pp2B*-14D-RNAi; *Dmef2*-*Gal4*), (D”’) *canA1* overexpression flies (*CanA1*-OE; *Dmef2*-*Gal4*) show normal myofibrillar structure.

### Overexpression of the constitutively active form of Calcineurin-A in IFMs affects the function but not the gross sarcomeric structure

A previous study has shown that overexpression of calcineurin in the heart results in cardiac hypertrophy (Molkentin et al., 1998). To check if overexpression of *canA1* in the IFMs leads to non-functional defective muscle, we over-expressed the *mcanA1*^*act*^ in developing *Drosophila* muscles using the *Dmef2*-Gal4 driver. This genetic combination was lethal to the progenies as very few adults were observed. The majority of the animals died at the early pupal stage and very few proceeded to the late developmental stage. Few flies that eclosed and survived, showed flightless phenotype (Fig. 1C), with no visible change in the structure of their IFMs at the fascicular level or the myofibrillar level (Fig. 1D”’) as compared to the control (Fig. 1D).

### Hopout mutants of *canB2* show defects in development

Since there are three isoforms of calcineurin in the IFMs, with similar structure and function, it was possible that they are functionally redundant. Therefore, in order to fully embrace the role of calcineurin in muscles, it was necessary to inactivate all the calcineurin isoforms in skeletal muscle. Since, it is practically difficult to reduce the levels of all three isoforms of Calcineurin-A together, we used another approach. It has been shown previously (Gajewski et al., 2003) that pupal developmental arrest induced by a forced expression of the constitutively active form of Calcineurin-A in the muscles of *Drosophila* is rescued by deleting the regulatory subunit of Calcineurin, *canB2*. This goes to show that the regulatory subunit of Calcineurin is essential for the functioning of all the catalytic subunits of the protein (Gajewski et al., 2003) and thus, the null of this gene will simultaneously reduce the activity of all the three Calcineurin A isoforms. A mutant allele of *canB2* was generated by hoping out P-element from the *canB2 EP(2)7074* parent line (Gajewski et al., 2003). Hopout line (referred as *EP(2)0774 Hopout*) generated was homozygous lethal at the early pupal stage. There were approximately 2% escapers observed for this genotype and all the escapers were flightless (Fig. 2A). The homozygous mutant escapers had severe defects in their muscle structure and showed degenerated fascicles (Fig. 2C,C’). Most of the muscle mass in these flies was localized to the either end of the thorax (Fig. 2C,C’). Myofibrils showed degenerating Z-discs and the sarcomere structure was lost in the majority of the myofibrils (Fig. 2C’). We were unable to detect any transcript of *canB2* in mutant flies by semiquantitative RT-PCR (Fig. 2D). In order to characterize the mutation in the *canB2*, both the regulatory region and the coding sequence was amplified using primers spanning the gene. We identified that region between promoter and the first exon is deleted (Fig. 2D’). These results implied that the allele, *EP(2)0774 Hopout* is a null for the *canB2*.

**Fig.2.**
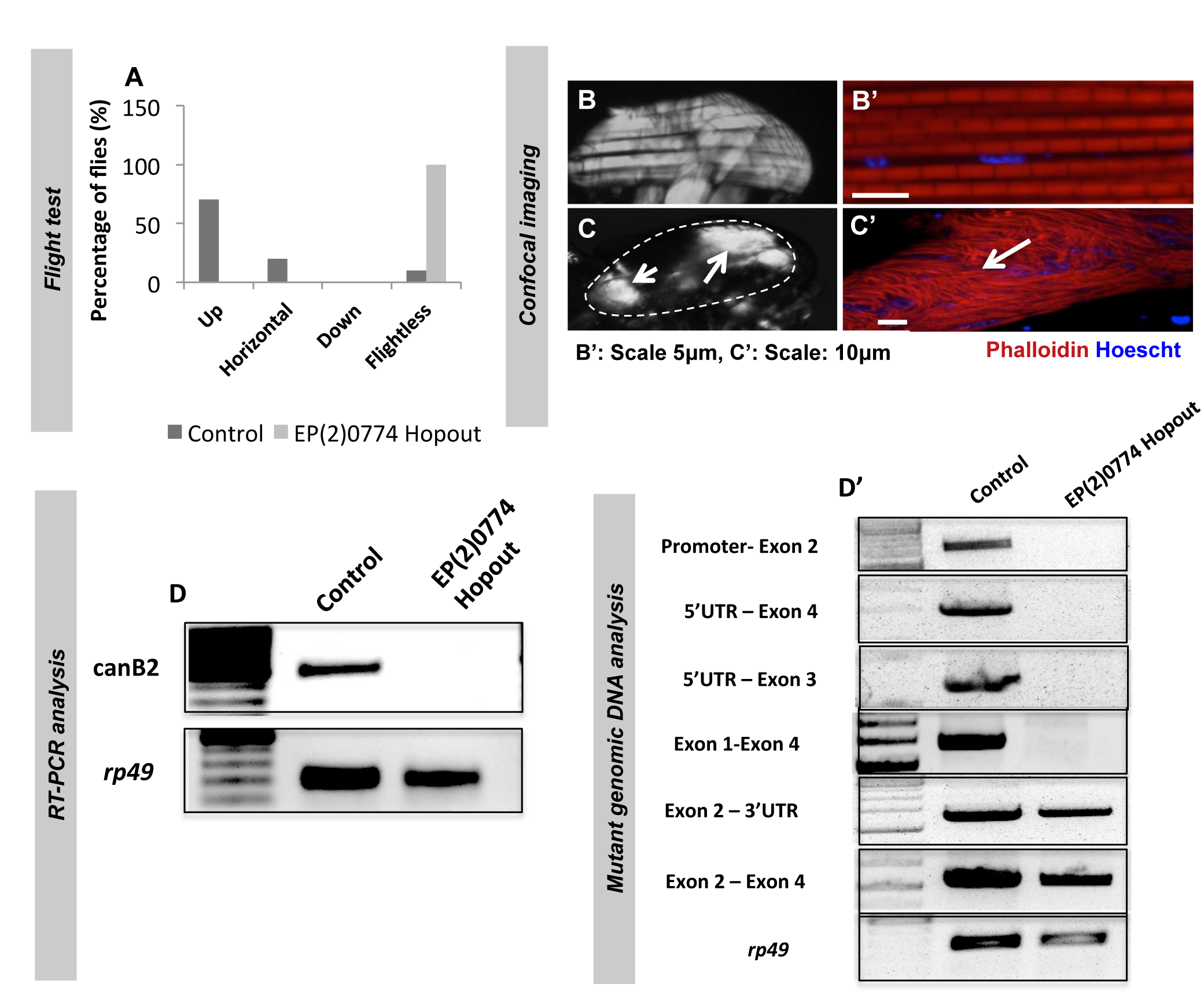
Muscle degeneration in adult *canB2* null flies. (A) Flight test shows that all the *canB2* mutant flies are flightless. (B and B’) Polarized and confocal imaging respectively of wild-type adult flies shows normal fascicle organization and myofibrillar structure. (C and C’) Polarized and confocal imaging respectively of homozygous *canB2* null mutants *(EP(2)0774 Hopout/EP(2)0774 hopout)* show disorganized fascicles and broken myofibrils. Arrows indicate muscle mass accumulated on the sides of thorax (D) RT-PCR shows expression of *canB2* in Wild-type (lane 1) and *canB2* null *(EP(2)0774 Hopout /EP(2)0774 Hopout)* (lane 2) animals. (D) Amplification of region of *canB2* gene from genomic DNA of wild-type and homozygous *canB2* null mutants *(EP(2)0774 Hopout/EP(2)0774 Hopout)* using primers flanking different exons shows absence of gene region between promoter and exon 2 of *canB2* gene (First panel: Promoter (Forward)-Exon2 (Reverse), Second panel: 5’UTR (Forward)- Exon 4 (Reverse), Third panel: 5’UTR (Forward)-Exon 3 (Reverse), Fourth panel: Exon 1 (Forward)- Exon 4 (Reverse), Fifth panel: Exon 2 (Forward)-3’UTR (Reverse), Sixth panel: Exon 2 (Forward)- Exon 4 (Reverse), Seventh panel: *rp49* (internal control)).

### Loss of *canB2* affects the IFM function and structure

In order to confirm that the defect seen in the *canB2* mutant is due to a loss of function of *canB2,* we specifically reduced levels of *canB2* in muscles by using Gal4-UAS system (Brand and Perrimon, 1993). *canB2* expression was reduced in muscles by using two different Gal4 lines. The *SG29.1*-Gal4 (Shyamala and Chopra, 1999) expresses in subsets of neurons and the IFMs, beginning early in the IFM development. *UH3*-Gal4 (Singh et al., 2014), on the other hand, expresses late in the development and becomes IFM-specific 36 hrs after puparium formation (APF). Using the Gal4-UAS system and *UAS-Dicer* (Dietzl et al., 2007) efficient knockdown of *canB2* was achieved in the IFMs (Fig. 3A,B). Flight test revealed that the majority of the flies are flightless when the knockdown is done with *SG29.1*-Gal4 (Fig. 3C). On the contrary, the majority of the flies were up-flighted when *canB2* was knocked down using *UH3*-Gal4 suggesting a temporal role of the protein in muscle development and function (Fig. 3C’). Flightless behavior in Drosophila can arise from the structural defect in the IFMs. In order to ascertain the cause of flightlessness in *canB2* deficient flies, structure of the IFMs was analyzed using polarized microscopy. There was a severe defect in the fascicular structure of the DLMs of flies in which *canB2* was reduced from the early developmental stages of the IFMs. Heterogeneity was observed in the phenotype owing to the penetrance effect and flies showed degeneration of muscles to various degrees (Fig. 3D,F’,F”). The number of flies showing muscle degeneration phenotype was around 60% following knockdown of *canB2* using *SG29.1-*Gal4 (Fig. 3H). On the contrary, the number of flies showing muscle degeneration (Fig. 3G’,G”) was significantly less (~10%) when *canB2* expression was reduced after 36 hrs APF (using *UH3*-Gal4) (Fig. 3I).

**Fig.3.**
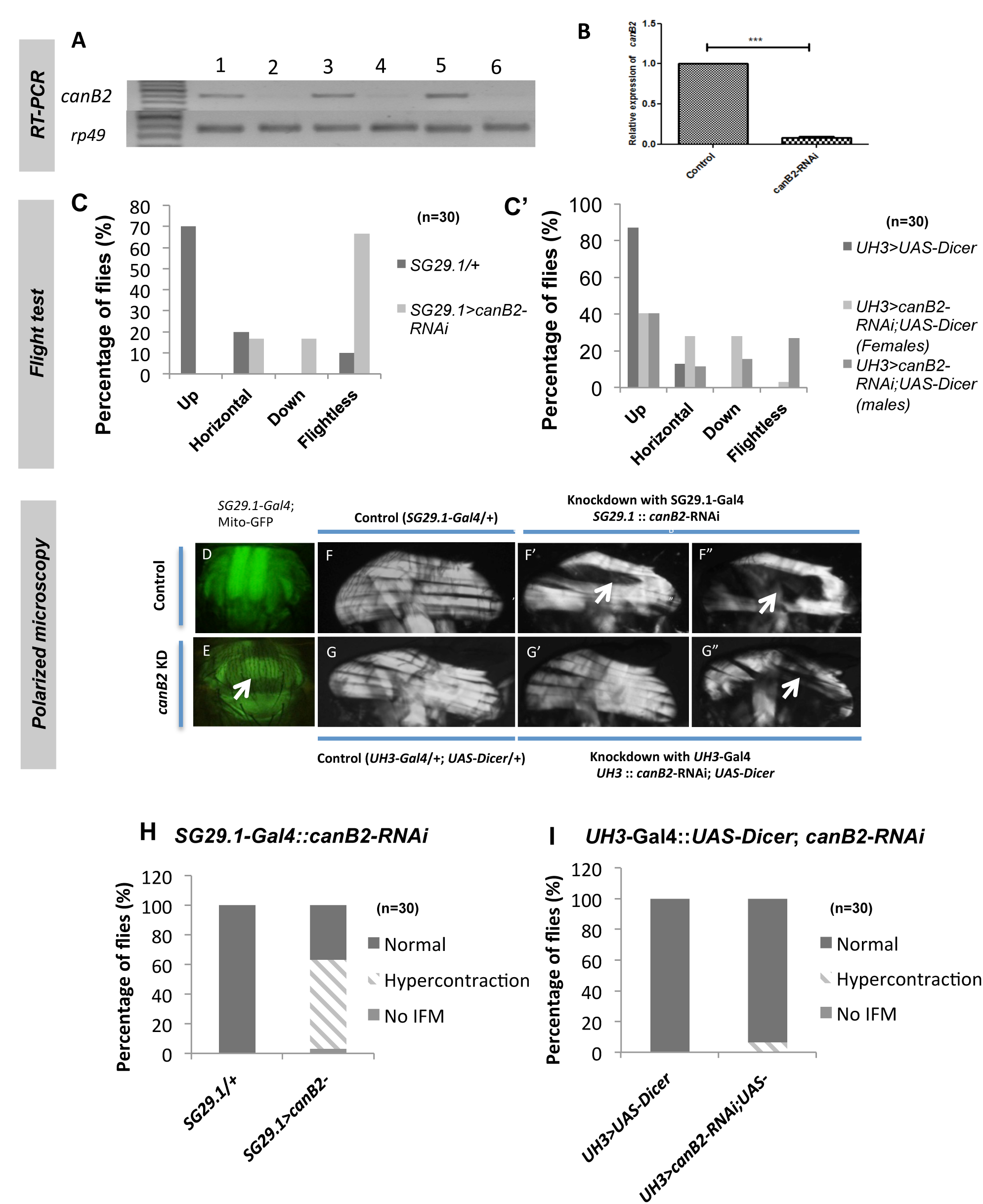
Depletion of *canB2* in muscles leads to muscle degeneration. (A) RT-PCR showing expression of *canB2* in wild-type and canB2 deficient background. (Lane 1: Ladder, lane 2: *SG29.1*-Gal4/+, Lane 3: *SG29.1*-Gal4; *canB2*-RNAi, Lane 4: *UH3*-Gal4/+, Lane 5: *UH3*-Gal4; *canB2*-RNAi, Lane 6: *UH3*-Gal4; *Gal80*^*ts*^; *canB2*-RNAi, Lane 7: *UH3*-Gal4; *Gal80*^*ts*^; *canB2*-RNAi). (B) Quantitative real time PCR showing expression of *canB2* in wild-type and *canB2* deficient adult flies as normalized to *rp49.* (C) Flight test shows that the majority of the flies with reduced levels of *canB2* (*SG29.1-*Gal4; *canB2*-RNAi) have defects in the flight (*SG29.1-*Gal4/+) (n=30) (C’) Flight test of flies in which levels of *canB2* are reduced post 36hrs APF (*UH3-Gal4*; *Gal80*^*ts*^ *UAS*-*Dicer*, *canB2*-RNAi) show no changes in the flight as compared to the control (*UH3-Gal4*; *Gal80*^*ts*^; *canB2*-RNAi) (D and E) IFMs in the adult Control (*SG29.1-Gal4*; *UAS*-mito-GFP) and *canB2* deficient (*SG29.1*-Gal4; mito-GFP; *canB2*-RNAi) flies respectively as marked by mito-GFP. Arrow indicates missing IFM fascicle in *canB2* deficient flies. (F) Polarized image of control (*SG29.1*-Gal4/+) shows six normal fascicles. (F’ and F’’) Representative polarized images of IFMs of adult flies with reduced levels of *canB2* (*SG29.1-Gal4*; *canB2*-RNAi). Arrows indicate degenerated muscle fascicles. (G) Polarized image of Control (*UH3*-Gal4; *Gal80*^*ts*^; *UAS*-*Dicer*). (G’ and G”) Representative polarized images of IFMs of adult flies with reduced levels of *canB2* (*UH3-Gal4*; *Gal80*^*ts*^; *UAS*-*Dicer*, *canB2*-RNAi). Arrow indicates degenerated fascicles. (H and I) Quantification of the number of flies showing muscle degeneration phenotype. (H) 60% of the flies with reduced levels of *canB2* (*SG29.1*-Gal4; *canB2*-RNAi) from the beginning of IFM development show degenerated muscles. (I) Knockdown of *canB2* post 36hrs APF (*UH3*-Gal4; *Gal80*^*ts*^; *UAS*-*Dicer*, *canB2*-RNAi) causes muscles degeneration in 10% of flies.

The sarcomere is the functional unit of the muscle that harbors an elaborate assembly of proteins required for proper contraction. Any defect in the contractile machinery or the structure of sarcomere leads to irregular contraction and muscle degeneration. The myofibrillar structure of the IFMs was analysed from both control and flies with knockdown of *canB2.* As compared to the control (Fig. S1A,A’), which had a normal organization of the sarcomeres, flies with knockdown of *canB2* had degenerated sarcomeres and broken Z-discs (Fig. S1B,B’). TEM images also revealed degenerated fibrils and mitophagy in the IFMs (supplementary material Fig. S1B”). In the transverse view of the IFMs, control showed normal 6 fascicles and close-packed arrangement of the myofibrils (Fig. S1C,C’) whereas a reduction in the expression of *canB2* led to the loss of fascicles and perturbed the close-packed arrangement of myofibrils (Fig. S1D,D’).

### Reduction in *canB2* expression in muscles affects late IFM development

The adult DLMs are the product of fusion of the myoblasts that have migrated from the proximal part of the wing disc (former VG-expressing adepithelial cells, Sudarsan et al., 2001) with the larval oblique muscles (LOMs, Fernandes et al., 1991). Following myoblast fusion and DLM splitting, the process of myofibrillogenesis begins around 30 hrs APF. Although we observed the muscle defect in the adult IFMs of *canB2* deficient flies, it is not clear if the defects seen in adult myofibrils were manifestation of the assembly deformity during developmental stages. In order to elucidate the importance of calcineurin in IFM development, canB2 was knocked down and the process of IFM formation was monitored from 0 hrs till 90 hrs APF. Control showed normal development of IFMs at all the stages observed (Fig. S1G,G’,G”,G”’). In spite of reduction in *canB2* expression, the dorso-longitudinal muscles (DLMs) are fully formed and mostly attached to the cuticle at around 50 hrs APF. Therefore, we conclude that knockdown of *canB2* causes no major defects of myoblast fusion, myotube formation, and fiber formation during IFM development. However, IFMs of *canB2* deficient animals start degenerating at around 60 hrs APF (Fig. S1H”) when the muscles begin contraction, suggesting the requirement of *canB2* in late development of the IFMs (Fig. S1H,H’,H”,H”’). Myofibrillogenesis was monitored in both control and *canB2* knockdown flies from 34 hrs till 60 hrs APF and there was no change visible in the sarcomere pattern of the myofibres of *canB2* deficient muscles (Fig. S1D,D’,D”,D”’), as compared to the control (Fig. S1I,I’,I”,I”’).

### Loss of *canB2* leads to muscle hypercontraction that is fully suppressed by muscle myosin heavy chain mutant, *Mhc*^*P401S*^

Irregular acto-myosin interaction can lead to excessive muscle contraction and subsequently contraction mediated muscle degeneration. Mutations causing IFM hypercontraction, such as *wupA*^*hdp-2*^ and *up*^*101*^, can be suppressed by genetically manipulating the amount of functional myosin assembled into thick filaments thereby countering the destructive unregulated muscle contraction (Nongthomba et al., 2003). *Mhc*^*P401S*^ is a myosin heavy chain mutant that has a defect in binding to F-actin. Replacing one *Mhc*^*+*^ gene copy with the *Mhc*^*P401S*^ mutant in *canB2* deficient flies (*SG29.1-*Gal4*/+; Mhc*^*P401S*^*/Mhc*^*+*^*; UAS-canB2-RNAi*) produces flies with normal wing posture and normal IFM morphology under polarized light (supplementary material Fig. S1F,F’). This suggests that the hypercontraction phenotype of *canB2* deficient flies involves irregular acto-myosin activity similar to mutants of troponin-I and troponin-T (Nongthomba et al., 2003; 2007).

### Genetic interaction between hypercontracting mutants of troponin Complex and canB2 reveals synergistic interaction between calcineurin and troponin-T

To further define how the loss of *canB2* affects IFM structure, we analyzed genetic interactions with mutants of Troponin-T and I that increase the contractile state of the muscle. Mutations in troponin-I, *wupA*^*hdp-2*^ and *wupA*^*hdp-3*^, and troponin-T, *up*^*1*^ and *up*^*101*^ have been previously identified as hypercontraction mutations (Nongthomba et al., 2003; 2004; 2007; Cammarato et al., 2004). *wupA*^*hdp-3*^ and *up*^*1*^ are nulls of troponin-I and troponin-T respectively and are categorized as stoichiometry mutants whereas *wupA*^*hdp-2*^ and *up*^*101*^ mutants have disturbed interactions with troponin-C and tropomyosin respectively and are therefore categorized as regulatory mutants (Nongthomba et al., 2004; 2007). To understand the interaction of calcineurin with structural proteins, *canB2* was knocked down in each of these mutant backgrounds using *SG29.1*-Gal4 and *UH3-*Gal4. A reduction in the expression of *canB2* in wild-type background during early muscle development resulted in hypercontraction phenotype in the majority of the animals. Knockdown of *canB2* in stoichiometry mutant background (*wupA*^*hdp-3*^ (Fig. 4B’,F) and *up*^*1*^ (Fig. 4C’,H)) did not result in either suppression or enhancement of the phenotype. On the contrary, knockdown of *canB2* in the regulatory mutant background (*wupA*^*hdp-2*^ (Fig. 4D’,G) and *up*^*101*^ (Fig. 4E’,I) resulted in enhancement of the phenotype with the majority of double mutant flies showing no muscles at all.

**Fig.4.**
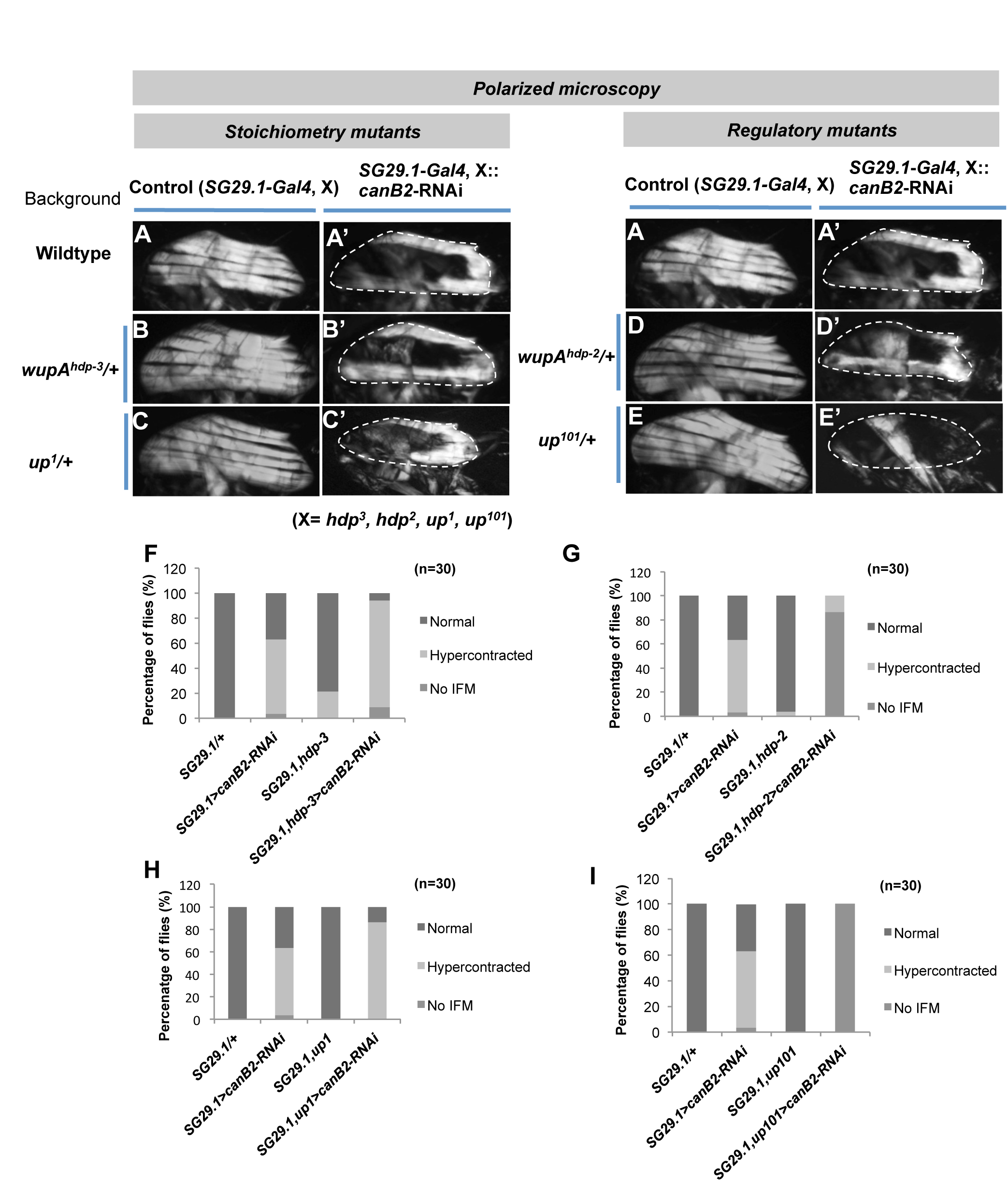
*canB2* shows genetic interaction with regulatory mutants of troponin I and T. (A, B, C, D, E) Polarized microscopy images of thorax of *Gal4* control adult flies in trans with mutants for Troponin I (*wupA*^*hdp-2*^*, wupA*^*hdp-3*^) and T (*up*^*1*^*, up*^*101*^) protein (A-*SG29.1*-*Gal4*/+, B-*SG29.1*-*Gal4*, *wupA*^*hdp-3*^, C-*SG29.1*-*Gal4*,*up*^*1*^, D-*SG29.1*-*Gal4*,*wupA*^*hdp-2*^, E-*SG29.1*-*Gal4*, *up*^*101*^). All the mutants are in heterozygous state and show normal organization of six fascicles in the thorax. (A’,B’,C’,D’,E’) Polarized microscopy images of hemithorax of *canB2* deficient flies in trans with mutants of Troponin I and T proteins (A’-*SG29.1*-*Gal4*/+; *canB2*-RNAi, B’-*SG29.1*-*Gal4*,*wupA*^*hdp-3*^; *canB2*-RNAi, C’-*SG29.1*-*Gal4*,*up*^*1*^; *canB2*-RNAi, D’- *SG29.1*-*Gal4*,*wupA*^*hdp-2*^*; canB2*-RNAi, E’- *SG29.1*-*Gal4*,*up*^*101*^; *canB2*-RNAi). *canB2* deficient flies in trans with *up*^*101*^ and *wupA*^*hdp-2*^ mutants show enhancement in the hypercontraction phenotype as opposed to *up*^*1*^ and *wupA*^*hdp-3*^ mutants. (F) Bar graph representation of the distribution of different muscle phenotypes (Normal, Hypercontracted, No IFM) of the following genotypes (From left to right, *SG29.1*-*Gal4*, *wupA*^*hdp-3*^*, SG29.1-Gal4*,*wupA*^*hdp-3*^; *canB2*-RNAi, *SG29.1*-*Gal4*/+, *SG29.1*-*Gal4*; *canB2*-RNAi) (G) Bar graph representation of the distribution of different muscle phenotypes (Normal, Hypercontracted, No IFM) of the following genotypes (From left to right, *SG29.1*-*Gal4*,*wupA*^*hdp-2*^*, SG29.1-Gal4*,*wupA*^*hdp-2*^; *canB2*-RNAi, *SG29.1*-*Gal4*/+, *SG29.1*-*Gal4*; *canB2*-RNAi) (H) Distribution of different muscle phenotypes as represented in the form of bar graphs for the following genotypes (From left to right, *SG29.1*-*Gal4*,*up*^*1*^*, SG29.1-Gal4*,*up*^*1*^; *canB2*-RNAi, *SG29.1*-*Gal4*/+, *SG29.1*-*Gal4*; *canB2*-RNA) (I) Distribution of different muscle phenotypes as represented in the form of bar graphs for the following genotypes (From left to right, *SG29.1*-*Gal4*, *up*^*101*^*, SG29.1-Gal4*,*up*^*101*^; *canB2*-RNAi, *SG29.1*-*Gal4*/+, *SG29.1*-*Gal4*; *canB2*-RNA)

Contrary to the above results, reducing the levels of *canB2* after the beginning of myofibrillogenesis in the wild-type background with *UH3-Gal4* affected the muscle structure of only 10% of flies (Fig. S2A’). Moreover, mutants of troponin proteins, *wupA*^*hdp-2*^ (Fig. S2C’,E) and *up*^*1*^ (Fig. S2B’,G) did not enhance the severity of hypercontraction phenotype observed for muscles deficient for *canB2*. However, knockdown of *canB2* in *up*^*101*^ (Fig. S2D’) background resulted in enhancement of phenotype and 100% of the double mutant flies showed hypercontraction phenotype (Fig. S2D’,F).

### *canB2* mutants show disturbed calcium homeostasis in the muscles

Muscular dystrophies are a group of hereditary diseases characterized by muscle fiber necrosis and progressive muscle wasting and weakness (reviewed in Campbell, 1995). Dystrophic muscles show excessive calcium influx into muscle fibres or disturbed intracellular calcium signaling which is presumably involved in the patho-physiology of muscle dystrophies (Jackson et al., 1985; Tutdibi et al., 1999). In order to understand the cause of muscle degeneration in *canB2* mutants, which show fibre necrosis and muscle degeneration like dystrophin mutants, calcium homeostasis was studied in the mutant muscles by expressing calcium sensor GCaMP. To determine whether the primary effect of the *canB2* null mutation is on muscle degeneration with consequent calcium homeostasis changes or vice versa, calcium imaging was done at 50hrs APF before muscles degeneration begins. As compared to the control (Fig. 5B,B’; Movie 1), calcium spikes in the mutant are broad (Fig. 5C,C’; Movie 3) suggesting a defect in calcium quenching ability of the system as visualized in the pseudocolour (Fig. 5A, control; 5A’, *canB2* mutant). The average peak area (Fig. 5E,E’, Table S1) and peak width (Fig. 5D,D’, Table S1) of calcium oscillations of *canB2* mutants was significantly higher than the control, suggesting that the amount of time free calcium spends in the cytosol or in the vicinity of myofibrils is higher in mutant as compared to the control.

**Fig.5.**
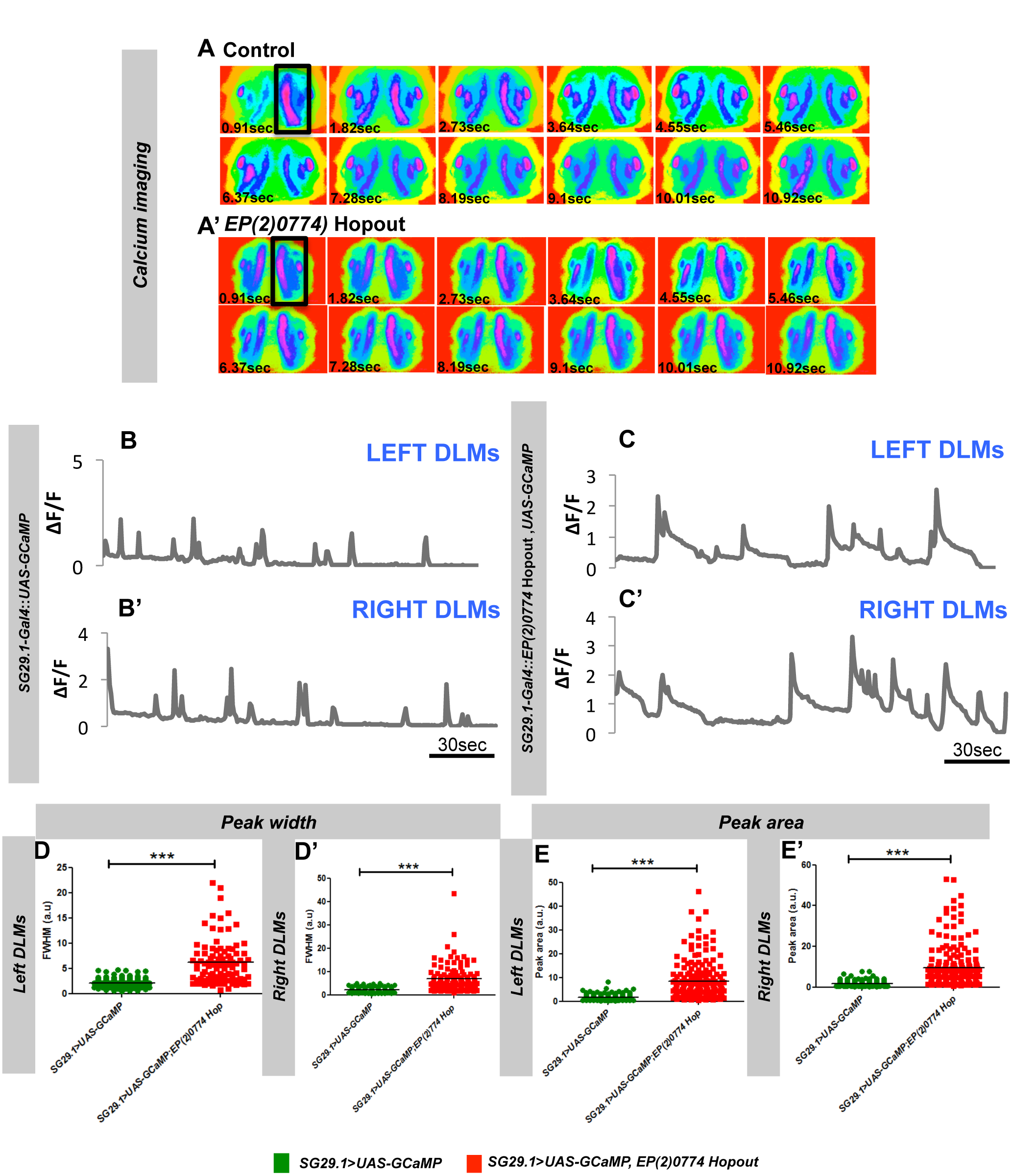
Impaired calcium homeostasis in indirect flight muscles of *canB2* null flies. (A) Pseudocolour images represent changes in cytosolic Ca^2+^ of indirect flight muscles in control and (A’) *EP(2)0774 Hopout* mutant IFMs over 11 secs as visualized by GCaMP3.0 calcium sensor. Recording was done at 1 frame/sec for 5 min. Only 12 frames are shown for one genotype, which indicates a randomly picked region from the calcium oscillation trace. Black box marks the region that shows change in the fluorescence. (B and B’) The traces are representative of spontaneous calcium oscillations in indirect flight muscles on the left (B) and the right side (B’) of the thorax of control 50hrs APF pupae (*SG29.1*-*Gal4*; *UAS*-*GCaMP*). (C and C’) The traces are representative of spontaneous calcium oscillations in indirect flight muscles on the left (C) and the right side (C’) of the thorax of *EP(2)0774 Hopout* mutant pupae (*SG29.1*-*Gal4*; *UAS*-*GCaMP; canB2-*RNAi). Recordings are representative of 18 (Control) and 7 (*EP(2)0774 Hopout* mutant) pupae. (D and D’) Peak width of spontaneous calcium spikes in DLMs from left (D) and right (D’) side of the thorax of *EP(2)0774 Hopout* mutant animals is significantly higher than the control (left DLMs P<0.0001, right DLMs <0.0001, Unpaired Student’s T test). (E and E’) Peak area of spontaneous calcium spikes in DLMs from left (E) and right (E’) side of the thorax of *EP(2)0774 Hopout* mutant animals is significantly higher than the control (left DLMs P<0.0001, right DLMs P<0.0001, Unpaired Student’s T test). Table S1 includes average values of peak width, peak area and peak frequency for above mentioned genotypes.

### Muscle-specific knockdown of *canB2* leads to disturbed calcium homeostasis in the muscles

Flight in Dipterans like *Drosophila* is controlled by both muscles (direct and indirect flight muscles) (Swank et al., 2006; Pringle, 1949) and neurons (giant fibre pathway) (Harcombe and Wyman, 1977; Koenig and Ikeda, 1980; Lehmann, 2016; Pathak et al., 2015; Tanouye and Wyman, 1980). IFMs are innervated by the motor neurons that fire at a much slower rate than the contraction frequency of muscles. They are necessary for maintaining myoplasmic [Ca^2+^] above the critical threshold that maintains the muscle in a stretch-activable state (Gordon and Dickinson, 2006). Since *canB2* is found at higher levels in both neurons and muscle it is conceivable that the changes observed in calcium spiking derive from a reduced *canB2* function in both these tissues. To test this, *canB2* was knocked down either in the muscles or in both neurons and muscles together and the calcium spiking activity was observed in the muscles.

#### a. Knockdown of *canB2* using neuronal and muscle-specific Gal4 lines

Knockdown of calcineurin was performed using *SG29.1-*Gal4 (Shyamala and Chopra, 1999), which expresses both in neurons and muscles. As compared to the control (Fig. S3B, left DLMs; B’, right DLMs; Movie 1), calcium spikes in muscles with reduced levels *canB2* are broader (Fig. S3C, left DLMs; C’, right DLMs; Movie 4) as visualized in the pseudocolour images over time (control, Fig. S3A; *canB2* knockdown, Fig. S3A’). Similar to the *canB2* mutants, the average peak area (Fig. S3D, left DLMs and D’, right DLMs; Table S1), peak width (Fig. S3E, left DLMs and E’, right DLMs; Table S1), peak frequency (Fig. S3F, left DLMs: F’, right DLMs; Table S1) of these oscillations in IFMs with reduced expression of *canB2* is significantly higher than that of the control.

#### b. Knockdown using muscle-specific Gal4

Calcineurin was knocked down in muscles alone using *Dmef2-*Gal4, which expresses specifically in the muscles. Calcium spiking in muscles with reduced expression of *canB2* (Fig. 6C, left DLMs; C’, right DLMs; Movie 5)) is different than control (Fig. 6B left DLMs; B’, right DLMs; Movie 2). Pseudocolour images show the changes in fluorescence in both control (Fig. 6A) and *canB2* knocked down muscles (Fig. 6A’) over period of 5 min. Similar to the *canB2* mutants, the average peak area (Fig. 6E, left DLMs; E’, right DLMs; Table S1), peak width (Fig. 6D, left DLMs; D’, right DLM; Table S1), peak frequency (Fig. 6F, left DLMs; F’, right DLMs; Table S1) of oscillations in *canB2* deficient IFMs was significantly higher than the control.

**Fig.6.**
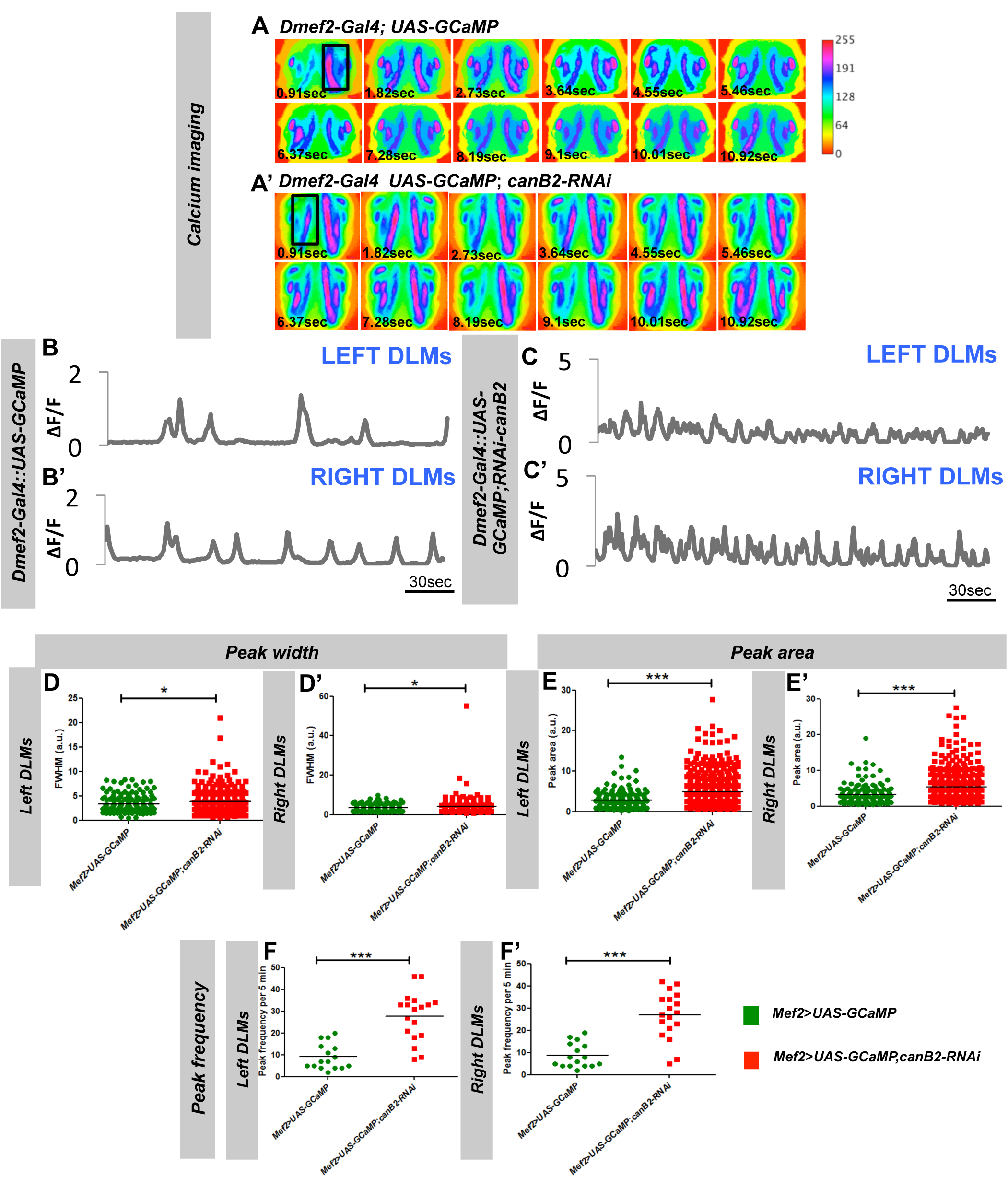
Impaired calcium homeostasis in indirect flight muscles of flies deficient for *canB2* in muscles alone. (A) Pseudocolour images representchanges in cytosolic Ca^2+^ of indirect flight muscles in control and (A’) *canB2* deficient (*UAS*-GCaMP; *canB2*-RNAi, *Dmef2*-*Gal4*) IFMs over 11 secs as visualized by GCaMP3.0 calcium sensor. Black box marks the region that shows change in the fluorescence. (B and B’) The traces are representative of spontaneous calcium oscillations in indirect flight muscles on the left (B) and the right side (B’) of the thorax of control 50hrs APF pupae (*UAS*-*GCaMP; Dmef2-Gal4*). (C and C’) The traces are representative of spontaneous calciumoscillations in indirect flight muscles on the left (C) and the right side (C’) of the thorax of *canB2* deficient pupae (*UAS*-*GCaMP; canB2-*RNAi, *Dmef2*-*Gal4*). Recordings are representative of 18 (Control) and 18 (*canB2-KD*) pupae. (D and D’) Peak width of spontaneous calcium spikes in DLMs from left (D) and right (D’) side of the thorax of *canB2* deficient pupae is significantly higher than the control (left DLMs P=0.0276, right DLMs P=0.0327, Unpaired Student’s T test). (E and E’) Peak area of spontaneous calcium spikes in DLM’s from left (E) and right (E’) side of the thorax of *canB2* deficient pupae is significantly higher than the control (Left DLMs P value= P<0.0001; Right DLMs P<0.0001), Unpaired Student’s T test). (F and F’) Peak frequency of calcium spikes in *canB2-KD* flies from both left (F) and right (F’) DLMs is significantly higher than control (left DLMs P<0.0001, right DLMs P<0.0001, Unpaired Student’s T test). Table S1 includes average values of peak width, peak area and peak frequency for above mentioned genotypes.

### Overexpression of the constitutively active form of Calcineurin-A suppresses calcium oscillations in IFMs

Previous results show that overexpression of constitutively active *canA1* does not affect the structure of IFMs (Fig. 1D”’). However, we observed defect in the function of these muscles. We therefore, checked for the muscle functioning by monitoring calcium homeostasis in this tissue. Calcium imaging of IFMs overexpressing *canA1* showed no calcium spikes at the time points measured (34 hrs APF, 42 hrs APF, 50 hrs APF and 60 hrs APF) (Data is shown only for 50 hrs APF) (Fig. S4C,C’). Control, on the other hand, showed normal spiking activity in muscles at 50 hrs APF (Fig. S4B,B’). The frequency of calcium oscillations in muscles with higher expression of *canA1* was zero (Fig. S4D,D’, Movie 6; Table S1). The absence of spiking, however, did not affect the development of the muscles, which suggests that myofibrillar assembly is not depended on spontaneous calcium oscillations seen in early developing IFMs. Thus, absence of the calcium oscillations in *canA1* overexpressing flies results in flightless phenotype.

### Increasing the levels of calcium in *canB2* knockdown background enhances the hypercontraction phenotype

As observed previously, *canB2* mutants show broad calcium spike suggesting that the amount of time free calcium remains in the cytosol is higher for muscles with reduced levels of calcineurin as compared to the wild-type. Hypercontraction in *canB2* deficient muscles can, therefore, result from increased and continuous exposure of myofibrils to calcium, which is otherwise controlled in wild-type muscles. Therefore, reducing the levels of calcium should decrease the muscle contraction and hence muscle degeneration. Considering the number of calcium channels in muscles and the complexity of their regulation, it is technically challenging to reduce the amount of calcium by decreasing the function of each of the channels simultaneously. However, instead of reducing the level of calcium in the muscle, we increased the calcium in the cytosol using mutant of *dSERCA* (*Kum170*), to see if the opposite holds true. *Kum170* is a loss-of-function mutant which has defect in either ATP binding or conformational state of the molecule (Sanyal et al., 2005). Increasing calcium level in *canB2* knockdown background genetically by using mutants of *dSERCA* (*Kumbhakarna*) had an adverse effect on the structure of muscles (Fig. 7D,D’). One copy of *Kum170* is sufficient to increase the severity of the hypercontraction phenotype observed for *canB2* mutants (Fig. 7D,D’,G). We also observed that the area of muscles in the thorax of flies, deficient for both *canB2* and *dSERCA*, was lesser than that of flies deficient for *canB2* only (Fig. 7H). Similarly, when the expression of *dSERCA* and *canB2* was simultaneously reduced, an enhancement in the hypercontraction phenotype was observed (Fig. 7E,E’,F). Area of indirect flight muscle in the double mutants (*SG29.1*-Gal4; *canB2*-RNAi, d*SERCA*-RNAi) (Fig. 7H) was found to be lesser than that of the control and *canB2* deficient flies alone.

**Fig.7.**
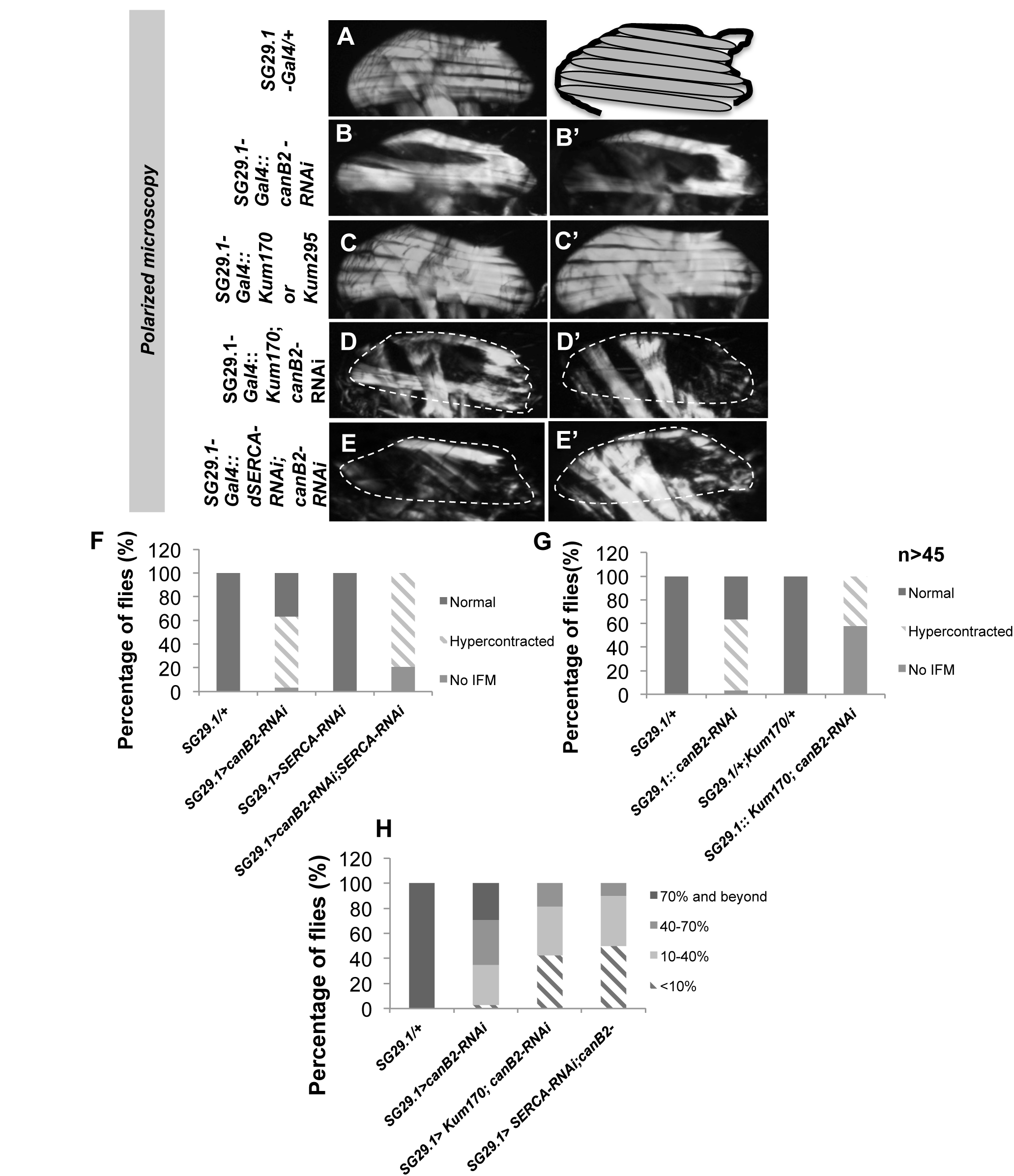
Increasing the level of calcium causes enhancement in the severity of the muscle phenotype in *canB2* deficient flies. (A) Polarized micrographs of*Gal4*control adult flies (*SG29.1*-*Gal4*/+) show normal fascicle organization. (B and B’) Polarized micrograph images of *canB2* deficient flies show hypercontracted muscles (*SG29.1*-*Gal4*; *canB2-RNAi*). (C and C’) Representative polarized micrographs of heterozygous mutant of *dSERCA*, *Kumbhakarna170 (Kum170)* shows normal fascicle organization and number. (D and D’) Representative polarized micrographs of *canB2* deficient flies in trans with heterozygous mutant of dSERCA, *Kum170* (*SG29.1*-*Gal4*; *Kum170*/+; *canB2-RNAi*). (E and E’) Polarized micrographs of flies deficient for both *canB2* and d*SERCA* show enhancement in the severity of muscle phenotype shown by *canB2* deficient flies. (F) Bar graph representation of distribution of muscle phenotype in flies deficient for both *canB2* and d*SERCA* (From left to right, Column 1: *SG29.1*-*Gal4*/+, Column 2: *SG29.1*-*Gal4*; *canB2*-*RNAi*, Column 3: *SG29.1*-*Gal4;* d*SERCA*-RNAi, Column 4: *SG29.1*-*Gal4*; *canB2*-RNAi, d*SERCA*-*RNAi*). (G) Bar graph representation of distribution of muscle phenotype in flies deficient for both *canB2* in trans with *Kum170* mutant (From left to right, Column 1: *SG29.1*-*Gal4*/+, Column 2: *SG29.1*-*Gal4*; *canB2*-RNAi, Column 3: *SG29.1*-*Gal4*; *Kum170/+*, Column 4: *SG29.1*-*Gal4*; *Kum170/+*; *canB2*-*RNAi*). (H) Bar graph representation of area of muscles in thorax of flies with following genotype (Column 1: *SG29.1*-*Gal4*/+, Column 2: *SG29.1*-*Gal4*; *canB2*-*RNAi*, Column 3: *SG29.1*-*Gal4*; *Kum170/+*; *canB2*-*RNAi*, Column 4: *SG29.1*-*Gal4*; *canB2*-*RNAi*, d*SERCA*-*RNAi*)

### Increasing the levels of calcium in *up*^*101*^ background enhances the hypercontraction phenotype

Human hypertrophic cardiomyopathies (HCM) include two dominant mutations I79N and R92Q that are located within the TnT-Tm interaction region (residues 70–180) and they affect the Ca^2+^ sensitivity in tension versus force measurements (Tobacman et al., 1999). Similar to human mutations, *up*^*101*^, a TnT mutation in Drosophila lies within the calcium sensitive region in TnT-Tm interaction and causes hypercontraction in muscles when present in homozygous form (Naimi et al., 2001; Nongthomba et al., 2007). The vast majority of the *up*^*101*^ thin filaments in the absence of Ca^2+^ were shown to be in the active contracting state (Viswanathan et al., 2014). Thus, the *up*^*101*^ mutant protein may alter the equilibrium of the Tm position at rest such that at any given time the majority of thin filament regulatory units are in the contracting state (Viswanathan et al., 2014). Thus, reducing the expression of *canB2* in the calcium sensitive *up*^*101*^ background may cause a drastic change in the structure of muscles by increasing the levels of calcium in the muscle. To test the possibility that the enhancement in the phenotype of calcium sensitive mutant *up*^*101*^ is indeed because of the increase in the levels of calcium, calcium concentration in the mutant muscles was genetically raised by using mutants of *dSERCA*. Flight test showed that both *up*^*101*^ and *Kum170* heterozygous flies are up-flighted whereas double mutants are majorly flightless or down flighted (Fig. 8A). Also at the structural level, the mutant, *Kum170* did not show muscle degeneration in heterozygous condition (Fig. 8C). 14% of *up*^*101*^ flies showed hypercontraction in muscles (Fig. 8B,E,E’,D). On the other hand, 35% of the double mutants showed hypercontraction phenotype (Fig. 8F,F’,D).

**Fig.8.**
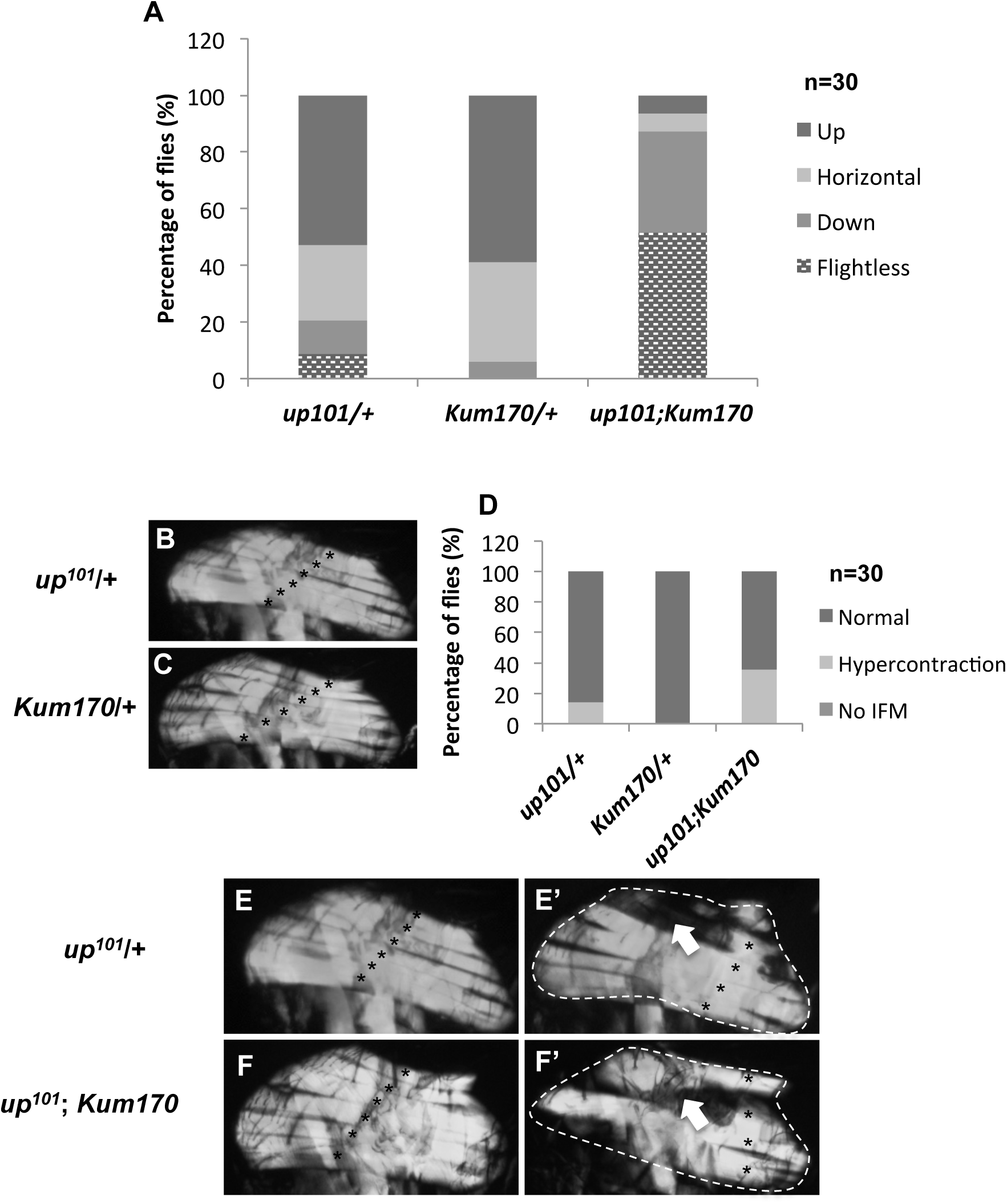
Increasing the level of calcium causes enhancement in the severity of the muscle phenotype of calcium sensitive *up*^*101*^ mutant. (A) Flight test shows that one copy of mutant protein (*troponin T*, *up*^*101*^ and *dSERCA*, *Kum170*) does not affect the flying ability of the adult flies. Double mutants (*up*^*101*;^*Kum170)* show flying defects and majority of flies are flightless. (B) Polarized micrograph images of hemithorax of flies heterozygous for *up*^*101*^ show normal fascicular organization. (C) Representative polarized micrographs of hemithorax of flies heterozygous for *Kumbhakarna170 (Kum170)* show normal fascicle organization and number. (D) Bargraph representation of the distribution of the muscle phenotype of flies heterozygous for *up*^*101*^, *Kum170* and double mutant of *up*^*101*^ and *Kum170*. Double mutants show more number of flies with hypercontracted muscles than single mutants alone. (E and E’) Polarized micrographs of DLMs of flies heterozygous for *up*^*101*^. Arrow indicate broken fascicles (F and F’) Polarized micrographs of DLMs of double mutants of *up*^*101*^ and *Kum170* show hypercontracted muscles. Arrow indicates degenerated fascicle.

## Discussion

### Calcineurin is involved in regulation of the spontaneous calcium activation during early myofibril assembly

It has been shown that spontaneous calcium transients play an important role during early myofibril assembly and is required for structural protein assembly into sarcomeres (Berchtold et al., 2000). We find that the IFMs also show calcium spikes as early as 34hr APF, when initial myofibrillar proteins are being laid to form sarcomeres (Reedy and Beall, 1993). The calcium spikes in the two adult IFM groups in each hemithorax ensure that they contract in a synchronized manner to share the mechanical load once muscle contraction is initiated (Gordon and Dickinson, 2006; Lehmann et al., 2013). However, in developing muscles, we find that the calcium spikes generated are cyclical (Movie 2) and they are not synchronized between the two IFM groups in each hemithorax. This suggests that these oscillations could arise spontaneously in the muscle without the neuronal cues. The calcium spikes are not rhythmic in our *canB2* knockdown or null pupae (Fig. 5C,C’; Fig. S3C,C’; Fig. 5C,C’) suggesting that calcineurin plays an important role in regulation of these spontaneous spikes. Calcineurin’s catalytic subunit has been previously shown to interact with calcium channels like ryanodine receptor (Bandyopadhyay et al., 2000) and voltage-gated calcium channels (Tandan et al., 2009) in vertebrates to regulate the flux of calcium from these channels. Disturbed calcium homeostasis seen in the IFMs with reduced levels of calcineurin suggests a role of calcineurin in the maintenance of calcium channels.

Our observation that the calcium spikes are completely abolished in the *canA*^act^ overexpressing lines and yet, the myofibrils assemble almost normally (Fig. 1D”’; Fig. S4C,C’) suggests that the spontaneous calcium transient spikes are dispensable for myofibril assembly in the IFMs. This phenomenon has been replicated by reducing free cytosolic calcium [Ca^2+^] with calcium buffering constructs and by down-regulating calcium channels in the IFMs (our unpublished data). This is also supported by our result that in *canB2* knockdown or null condition with arrhythmic calcium spikes, early myofibril assembly takes place normally (Fig. S1J,J’,J”,J”’) and the muscle abnormalities initiate only during later stages of myofibrillar growth (Fig. S1H”,H”’). So, unlike vertebrate embryonic (Ferrari et al., 1996) and myotube culture (Flucher and Andrews, 1993), spontaneous calcium activation is dispensable for myofibril assembly in the IFMs. But, as expected the calcium activation is required for muscle function (contraction) as all these genotypes show the flightless phenotype.

### Perturbed calcium response precedes development of hypercontraction muscle phenotype

Our study shows that reduction in the levels of *canB2* has a profound effect on the muscle structure with majority of the fibres showing degenerated Z-bands, broken or missing M-lines (indicative of reduced or missing thick filaments), and shorter sarcomeres (indicative of excessive contraction) (Fig. 2C,C’; Fig. S1B’,B”). Such phenotypes are also observed in other hypercontracting alleles (Nongthomba et al., 2003). Since the muscle phenotypes of hypercontracting alleles can be suppressed by reducing acto-myosin force, it can be concluded that assembly of the structural proteins takes place normally in these mutants. However, sustained strain as a result of unregulated acto-myosin interaction could cause muscle abnormalities in later stages of muscle development. This suggests that the interaction between the thick and thin filaments during early muscle development is in a controlled manner and uncontrolled acto-myosin interactions over time causes hypercontraction in the *canB2* mutant.

Our study demonstrates that the hypercontraction phenotype manifested by a reduction in the levels of CanB2 protein could be due to perturbed calcium homeostasis in the muscles and abnormalities in spontaneous calcium spikes (Fig. 5C,C’). Our data also strongly support the possibility that the production of arrhythmic calcium spikes precedes the development of microscopically noticeable abnormal muscle phenotype. The changes in calcium spikes that we have observed are also reminiscent of the defects in calcium sequestration observed for mutants of *dSERCA* (*Kum170*) in neuromuscular junctions (Sanyal et al., 2006). This essentially means that the period for which calcium remains in the cytosol is higher for *canB2* mutants thereby leading to uncontrolled contraction and untimely activation of several calcium-activated proteases like calpains leading to cell death. Possible substrates of calpains are the membrane cytoskeleton, the Ca^2+^-ATPase of the plasma membrane, and ion channel proteins (Salamino et al., 1994). The action of calpain on plasma membrane Ca^2+^channels can lead to increase openings of these channels (Turner et al., 1993). This could mean a positive feedback loop of Ca^2+^influx, Ca^2+^- dependent proteolysis, and increased Ca^2+^influx. The muscle degeneration phenotype seen in the canB2 mutants (Fig. 3F’,F”,G’,G”) could involve this pathway. Therefore, it is pertinent to suggest that calcium dysregulation may serve as diagnostic marker much before actual muscle abnormalities start to show up in any cardiomyopathies or/and other muscle disorders.

### Muscle hypercontraction results from increased calcium sensitivity of the myofibrils and unregulated acto-myosin interactions

The slow, progressive degeneration of IFMs in *canB2* mutants is similar to the hypercontracted IFM phenotype observed in mutants of myosin heavy chain (Kronert et al., 1995), troponin-T and troponin-I, (Kronert et al., 1999, Nongthomba et al., 2003, 2007) and flapwing (Pronovost et al., 2013). Unlike, troponin-I mutant, *wupA*^*hdp-3*^, that shows muscle defect as early as 50 hrs APF, there was no visible difference in the structure of the sarcomere in *canB2* knockdown flies at this stage. This suggests that the phenotype manifested in *canB2* knockdown flies is not a result of a reduction in levels of structural proteins rather it indicates defect in the regulation of structural proteins. We also confirmed this observation by checking levels of structural proteins in *canB2* deficient IFMs and we did not observe changes in the levels of transcripts of structural proteins like troponin-I, troponin-T, actin and tropomyosin (data not shown).

The troponin complex is a Ca^2+^-sensitive molecular switch that regulates acto-myosin interaction. The activity of the complex is modulated by phosphorylation of Troponin-T and Troponin-I proteins (Robertson et al., 1982; Solaro et al., 1976; Zakhary et al., 1999). *up*^*101*^, a mutant of troponin-T, harbors a mutation in troponin T-tropomyosin interacting region which enhances the calcium sensitivity of the troponin complex (Viswanathan et al., 2014). We found that raising the levels of calcium in the *up*^*101*^ background enhanced the severity of muscle degeneration phenotype (Fig. 8F,F’). Also, reduction in the levels of *canB2* in *up*^*101*^ heterozygous mutant background enhanced the severity of IFM defects of *canB2* mutant and the majority of flies had empty thoraces (Fig. 4E,E’; supplementary material Fig. S2D,D’). Since raising the level of calcium in muscles has similar effects on the enhancement of hypercontraction as reducing the level of *canB2* in the heterozygous *up*^*101*^ mutant, we believe that calcineurin is required for maintaining calcium homeostasis in the muscles. The enhancement of hypercontraction in the *up*^*101*^ mutant background (Fig. 4E,E’; Fig. S2D,D’) could be through buffering of calcium by troponin complex as explained by Montana and Littleton (2004). In wild-type muscles, calcium levels increase in the sarcomere following neuronal stimulation, in order to increase the fraction of troponin complexes active at a given point of time during regulated contraction. However, in mutants such as *up*^*101*^, calcium rather than returning to intracellular stores, remains buffered in the sarcomere. These mutations respond to lower calcium concentrations due to which the calcium remains bound to the troponin complex long after completion of a cycle of contraction. Another possibility suggested by Montana and Littleton (2004), reflects the ability of the *up*^*101*^ mutation to lower the activation energy of the troponin complex for the transition from inhibitory state to an activated state in the absence of calcium. Troponin complex in the activated state has a higher affinity for calcium that allows binding of calcium away from the endogenous muscle calcium buffers like calmodulin. This can also lead to an overall aberrant buffering of calcium. This condition is similar to human hypertrophic cardiomyopathy wherein the muscles are sensitized towards greater calcium adherence thereby prolonging mechanical relaxation and delaying calcium reuptake into SER (Davis et al., 2016). Taken together, increase in the free calcium [Ca^2+^] in the cytosol due to reduced expression of *canB2* and the overall buffering of calcium in the sarcomere of *up*^*101*^ mutants by the troponin complex enhances the severity of the hypercontraction phenotype. Supporting the above results, genetically raising the levels of calcium in the *canB2* knockdown background using mutants of *dSERCA* (*Kum170*), enhances the severity of hypercontraction phenotype (Fig. 7D,D’,E,E’). Heterozygous *dSERCA* mutants, on the other hand, do not show hypercontraction phenotype (Fig. 7C,C’) thereby reflecting on the diversity of pathways that calcineurin might affect, independent of the calcium homeostasis maintenance in the muscles. Isolated myofibrils and *in vitro* expression studies of many structural protein mutations responsible for cardiomyopathies and disorders showed sensitive calcium response (Reviewed in Tardiff, 2011; Harada and Potter, 2004; James et al., 2000). Our finding on the *up*^*101*^ mutation that is comparable to calcium sensitive TnT cardiomyopathies (Kooij et al., 2016) suggests that many of these mutations will show exacerbated phenotype with deregulated calcium homeostasis.

## Experimental Procedures

### Fly work

*Canton-S* was used as the wild-type control unless specified. Details of fly lines used in the study have been included in the supplementary. All the flies were maintained at 23±2°C, under 12 hrs Light/Dark (L/D) cycles on cornmeal-sucrose-yeast agar media. All Gal4-UAS experiments were done at 29°C unless specified.

### Flight test

The flight test was performed as described by Drummond et al., 1991. Briefly, flies were released into a Sparrow box, a plexiglass container with a light source at the top, to determine their ability to fly up, horizontal, down, or the inability to fly, denoted as flightless.

### Polarized light microscopy

Fly hemithoraces were prepared for polarized microscopy as described in Nongthomba and Ramchandra, 1999. Fly thoraces were frozen in liquid nitrogen and dissected from the midline. Thoraces were subsequently dehydrated in alcohol series and cleared using methyl salicylate and mounted using DPX medium. The hemithoraces were observed in Olympus SZX12 microscope under polarized light optics and photographed using Olympus C-5060 camera.

### Confocal microscopy

Flies were bisected from middle either using a blade or fine scissors in 4% Paraformaldehyde solution. Fixation of the tissue was followed by washing with phosphate buffer saline with 0.3% Triton-X100 (PBTx) for 1 hr (x4, 15 min). Samples were then stained using 1:40 diluted Phalloidin-TRITC (P1951, Sigma- 50ug/ml stock) for 1 hr and washed subsequently with PBTx for 30 min. Samples were mounted on a glass slide in Vectashield mounting media (Vector Laboratories, USA). Slides were then visualized under Zeiss 880 confocal microscope at 63X magnification with 543 nm wavelength.

### Transmission electron microscopy

Fly half thoraces were prepared following the protocol of Kronert et al., 1995. Adult hemithoraces were dissected in fixative (3% Paraformaldehyde, 2% Glutaraldehyde, 10 mM Sucrose, 100 mM Sodium phosphate pH 7.2 and 2 mM EGTA) (PBS, pH 7.2) and fixed overnight at 4°C in fixative. Tissues were washed with sodium phosphate buffer pH 7.2 (x2, 15 min), fixed with 1% osmium tetroxide in PBS buffer (30 min), washed with water for 15 min twice, block stained in uranyl acetate at room temperature for 1hr, washed with water (x2, 15min) and dehydrated through alcohol series, followed by final dehydration in absolute ethanol (x2, 30 min). Tissues were cleared using propylene oxide (x2, 15 min) and infiltrated with 1:1 propylene oxide: Epoxy Resin (overnight), followed by three changes (3x3 hours) in the embedding medium. Sections were cut (Leica EM UC6), stained with uranyl acetate for 2 hours, dried and stained with lead citrate for 5-7 min. Images were captured using a Tecnai G2 Spirit BioTWIN transmission electron microscope (FEI, Netherlands).

### Genomic DNA isolation

30 anesthetized flies were collected in an Eppendorf tube and frozen at -80°C. They were then ground in 400 µl Buffer A (100 mM Tris-HCl, pH7.5, 100 mM EDTA, 100 mM NaCl, 0.5% SDS) and incubated at 65°C for 30 min. After incubation, 800 µl of LiCl/KAc solution (1 part 5 M KAc stock: 2.5 parts 6 M LiCl stock) was added and incubated on ice for 10 min. The solution was spun and the supernatant was transferred into a new tube. 600 µl of isopropanol was added to the supernatant and mixed. Tubes were spun and pellet was washed with 70% alcohol and suspended in 150 µl TE. Primers used for gene amplification have been listed in supplementary information.

### RNA isolation and PCR

IFMs were dissected from the flies of the desired genotype and were kept in Trizol reagent till required. Total RNA was isolated using Trizol reagent (SIGMA) and RNA was quantified. 2 µg of the total RNA was used to make complimentary DNA (cDNA) using first strand cDNA synthesis kit (Maxima H minus First Strand cDNA Kit, Thermo scientific). 1µl of 1:10 diluted stock cDNA (2µg) was used in each reaction of semi-quantitative and Real-time PCR. For quantitative PCR, all reactions were carried out using SYBR green (DyNAmo SYBR Green qPCR Kit/F-410L, Thermo Scientific). Fluorescence intensities were recorded and analyzed in an ABI prism 7900HT sequence detection system (Applied Biosystems SDS 2.1 software, USA). The relative expression level was normalized using *rp49* as reference gene. The primer sequences have been listed in Supplementary information.

### Calcium imaging

For calcium imaging, GCaMP calcium sensor was expressed in the IFMs of the desired genotype using the Gal4-UAS system. Pupae of the desired genotype and time point were collected on a tissue paper. The pupal case of the pupae was removed and pupae were stuck to the glass slide. All recordings were done for 5 min using Olympus IX81 fluorescence microscope. All the images were analyzed using Image J (https://imagej.nih.gov/ij/). Quantification of the fluorescence signals was carried out by measuring the mean pixel value of an area in the IFMs for each frame of the movie using Image J (Version Image J 1.50i, Java 1.6.0_65 (64-bit). We defined the fluorescence change over time as ΔF/F = (F_0_ – F_t_)/(F_0_ - B), expressed in %, where F_0_ is the average pixel value in the first four frames of the experiment, F_t_ the fluorescence at any given time point, and B the average background intensity in an area without any muscle. Peak width and peak area value for each observed calcium peak was measured by OriginPro 8.5 software. Peak area is the area under the curve and signifies overall time taken by the peak to rise and fall. Peak width, on the other hand, is the width at the half maximum of the peak. Data is reported as average SEM.

### Statistical analysis

Data was plotted and statistical analyses were performed using GraphPad Prism 5 (GraphPad) and Microsoft Excel 2007. Unpaired student’s T-test was used to analyze all the graphs for statistical significance.

## Acknowledgement

We sincerely thank Prof. N. Gayathri at NIMHANS, Bangalore, for help with electron microscopy and subsequent discussions. We thank Mrs. Hemavathi for technical support during electron microscopy and Ms. Deepti Bapat, Ms. Divya at IISc Confocal Facility for their patience and interest. We extend our gratitude to Prof. Patrick O’Farrell (University of California, San Francisco) for the *UAS-canA*^*act*^ fly line. We thank the Bloomington Drosophila Stock Center, Vienna Drosophila Research Centre and the NCBS Stock Centre, Bangalore, India for providing flies.

## Author contributions

R.J. and U.N. developed the concept and designed the experiments. R.J. executed the experiments. R.J. and U.N. interpreted the data and wrote the manuscript.

